# Sent to the Corner: xylem vessel anatomy not surface area determines megaphyll hydraulics in *Cecropia obtusa* Trécul (Urticaceae)

**DOI:** 10.1101/259036

**Authors:** Sébastien Levionnois, Sabrina Coste, Eric Nicolini, Clément Stahl, Hélène Morel, Patrick Heuret

**Affiliations:** CNRS, UMR EcoFoG (AgroParisTech, CIRAD, CNRS, INRA, UA, UG), 97379 Kourou Cedex France.; UG, UMR EcoFoG (AgroParisTech, CIRAD, CNRS, INRA, UA, UG), 97379 Kourou Cedex France.; CIRAD, UMR AMAP (CIRAD, CNRS, INRA, IRD, Université Montpellier II), 34398 Montpellier, France.; INRA, UMR EcoFoG (AgroParisTech, CIRAD, CNRS, INRA, UA, UG), 97379 Kourou Cedex France.; CIRAD, UMR EcoFoG (AgroParisTech, CIRAD, CNRS, INRA, UA, UG), 97379 Kourou, France.

**Keywords:** allometry, *Cecropia obtusa*, Corner’s rules, hydraulic, intraspecific, leaf-stem relationship, petiole anatomy, lamina-petiole relationship

## Abstract

Corners rule predicts a positive correlation between leaf dimensions and the cross-sectional area of the primary stem. Although this relationship is usually explained by hydraulic and mechanical requirements, these hypotheses have never been tested empirically. However, Corner’s rule is tricky to investigate since rapid secondary growth of the stem prevents a rigorous link being established between a given leaf and the supporting stem. We chose a twig-like leaf model since petiole anatomy is only linked to the attached lamina. We tested the hypothesis that anatomical adjustments to hydraulic requirements related to vessel size enable reduced investment in tissue in the framework of Corner’s rule. We conducted a functional, mechanistic and intraspecific investigation of *Cecropia obtusa* Trécul, a Neotropical pioneer tree, by integrating morphological, anatomical and theoretical hydraulic traits around the lamina-petiole size relationship. The twig-like structure of the leaf and the strong lamina-petiole correlation of this model tree species made it possible to use the leaf-level model for a rigorous investigation of the functional implications of Corner’s rule. We found a positive correlation between petiole size, lamina size, the ratio of mean vessel area to vessel frequency in the petiole xylem and theoretical specific conductivity in the petiole xylem. Hydraulic function supports Corner’s rule to a lesser extent than previously thought. Variations in vessel dimensions mainly drive xylem hydraulic performances and avoid disproportionate petiole cross-sections to answer to hydraulic requirements associated with lamina size.

## INTRODUCTION

Leaves have received a great deal of attention because of the major role they play in the plant carbon economy thanks to photosynthesis. At the interspecific level, the currently widely accepted paradigm relates to a global spectrum of leaf economy, running from quick to slow returns on investments (Wright et al. 2004). Leaf surface area encompasses six orders of magnitude (Niinemets et al. 2007, Milla and Reich 2007, Wright et al. 2017). One possible and pertinent way for improving our understanding of such variability is to integrate the stem and to look at the size and functional relationship between stem and leaf.

From an evolutionary perspective, in his seminal paper on “Durian Theory” E.J.H. Corner already made empirical observations on such relationship, as “The stouter the main stem, the bigger the leaves and the more complicated their form” (Corner 1949). He completed and extended this with a second point he called “diminution on ramification” as “The greater the ramification, the smaller become the branches and their appendages”. These principles were referred to as Corner’s rules by Hallé *et al.* (Hallé et al. 1978) and have been explored by several authors (White 1983a, 1983b, Brouat et al. 1998, Brouat and McKey 2001, Westoby and Wright 2003, Preston and Ackerly 2003, Sun et al. 2006, Normand et al. 2008, Liu et al. 2010). However, from a more developmental point of view, even before Corner, by drawing attention to the pith, E.W. Sinnott (Sinnott 1921) suggested that the size of organs (the stem, leaf, fruit) is correlated with the size of the meristem from which they develop. Back to the contemporary research, plenty of studies have investigated both stem and leaf in their particular size and functional relationships (White 1983a, 1983b, Brouat et al. 1998, Ackerly and Donoghue 1998, Brouat and McKey 2001, Westoby and Wright 2003, Preston and Ackerly 2003, Sun et al. 2006, Kleiman and Aarssen 2007, Normand et al. 2008, Yang et al. 2008, 2008, Tao et al. 2009, Olson et al. 2009, Milla 2009, Liu et al. 2010, Whitman and Aarssen 2010, Scott and Aarssen 2012, Dombroskie and Aarssen 2012, Osada et al. 2015, Huang et al. 2016, Fan et al. 2017), usually (but not always) referring to Corner’s rule. Since a unifying framework is lacking, and based on the Corner’s vision, we will refer in this study to the leaf-stem relationship (LSR), that we defined as (i) the spectrum of size and functional variation in the relationship, (ii) between a leaf and its associated internode, or between leaves and the stem of a first-year shoot, and (iii) across or within species.

Both at the intra- and interspecific level, the LSR is frequently and intuitively explained by functional requirements in terms of hydraulic and mechanical supplies (Hallé et al. 1978, White 1983a, 1983b, Brouat et al. 1998, Westoby and Wright 2003, Normand et al. 2008). Since the respective contributions of the hydraulic and mechanical functions have not yet been tested, there is a need to (i) explore the diversity of anatomical structures that lead to interspecific variation in the LSR and (ii) understand the anatomical adjustments that allow for variations in the size of the stem and the leaf at the intraspecific level (Lehnebach et al.). The aim of this study was thus to investigate the intraspecific mechanisms of the LSR from the point of view of hydraulics and the aforementioned carbon allocation issues.

In angiosperms, xylem vessels drive water conduction through the soil-plant-atmosphere continuum (Tyree and Zimmermann 2002) according to the cohesion-tension theory (Dixon and Joly 1895, Steudle 2001). Vessel dimensions vary considerably at interspecific level (Chave et al. 2009, Zanne et al. 2010, Olson and Rosell 2013), at intraspecific level (Lens et al. 2011, Hajek et al. 2016) and at the intra-individual level (Anfodillo et al. 2006, 2013, Bettiati et al. 2012, Schuldt et al. 2013), with critical consequences for hydraulic performance at the organ, individual, population or species levels (Hacke et al. 2016). According to the Hagen-Poiseuille law (Tyree and Zimmermann 2002), vessel hydraulic conductivity is a fourth-power function of the vessel diameter. Thus, even slight variations in vessel diameter have dramatic impacts on associated conductance. A greater or lesser water demand with respect to the leaf lamina surface can theoretically be satisfied either by increasing the surface area of the conductive tissues which supply it, or by modifying the conductivity of the related tissues by means of their anatomy. Therefore, we hypothesise that adjustments to stem vessel size first drive size-related hydraulic supplies, thereby limiting carbon increments of xylem surface with an increase in leaf size. Consequently, even if stem xylem surface area is positively correlated with leaf size, the hydraulic function explains the LSR to a lesser extent than previously thought.

However, focusing on the leaf-stem system is hampered by several technical limitations. One of the main difficulties in applying a rigorous approach to the LSR is defining where to sample the last elongated shoot. On one hand, Corner (Corner 1949) refers to primary diameter (Hallé et al., 1978), which means that no secondary growth has occurred. Secondary growth occurs very soon, possibly at the same time as the elongation process, meaning that LSR may vary over time. On the other hand, the internode of a first-year shoot supplies not only the leaf at its apex, but all the leaves supported by internodes distal to it as well as the bud content (Cochard et al. 2005). It would thus be more relevant to analyse the relationship between the total surface area of leaves borne by the first-year shoot and the cross-sectional area of the shoot (Brouat et al. 1998). A compromise for a rigorous functional assessment would be to shift to a leaf-level model (i.e. considering the petiole to be homologous to the stem and its associated lamina), since there are plant groups in which the differences between leaf and shoot, or more precisely between the petiole and the stem remain fuzzy (Givnish 1984, Poorter and Rozendaal 2008, Nicotra et al. 2011). Indeed, compound or double-compound leaves, or large palmate or palmatilobate leaves look like more to a branch in a physiognomic or functional sense. For these species, the whole bifurcation structure of the tree for foraging space is realised by leaves and stems together. A major symptom of the transfer of the bifurcation function from stems to leaves is less stem branching intensity in comparison with simple leaved tree species (Aarssen 2012). Developmental genetics approaches also strengthen our posture since numerous studies suggest that the petiole should be considered separately from the leaf lamina, from morphogenesis and developmental genetic points of view, and more as an stem-like axial organ (Foster 1936, Tsukaya et al. 1993, 1995, 2002, Tsukaya 1995, 2013, Busch et al. 2011).

Using a leaf model to investigate the LSR makes it possible to avoid the bias frequently encountered at the phytomer level. Indeed (i) a petiole strictly supports mechanical and hydraulic constraints associated with its single lamina since it represents the closest structure to the transpiring compartments (Bucci et al. 2003) and (ii) even if a petiole undergoes secondary growth, the growth still only concerns the functional requirements of the associated lamina. Consequently, using a leaf model would make it possible to focus on petiole or venation traits, for instance, and to rigorously investigate the functional adjustments underlying the LSR. Finally, the lamina-petiole relationship has already been proven to be a relevant tool for functional investigations, and, for instance, (Royer et al. 2007) showed that this relationship is a significant predictor for inferring fossil functional traits.

In this context, the genus *Cecropia* (Urticaceae), an iconic pioneer tree in the Neotropics, could provide several consistent tree models to improve our understanding of structure-function relationships. These plants have a simple architecture (Heuret et al. 2002, Zalamea et al. 2008) with few botanical entities (e.g. axes, leaves), but have large petioles, reaching more than 1 m in length and up to 2 cm in diameter (Fig. 1b). Linked to its heliophile habit, petioles have a somewhat branching function (Givnish 1984, Poorter and Rozendaal 2008) and can be considered as axes, since the hydraulic and mechanical features of petioles are comparable to a branch with respect to the form of the leaf (palmatilobate to palmate megaphylls with numerous lobes tens of centimetres in length) and its size (up to 0.4 m^2^ for *Cecropia obtusa* Trécul with 1-m long petiole).

**Fig 1.**
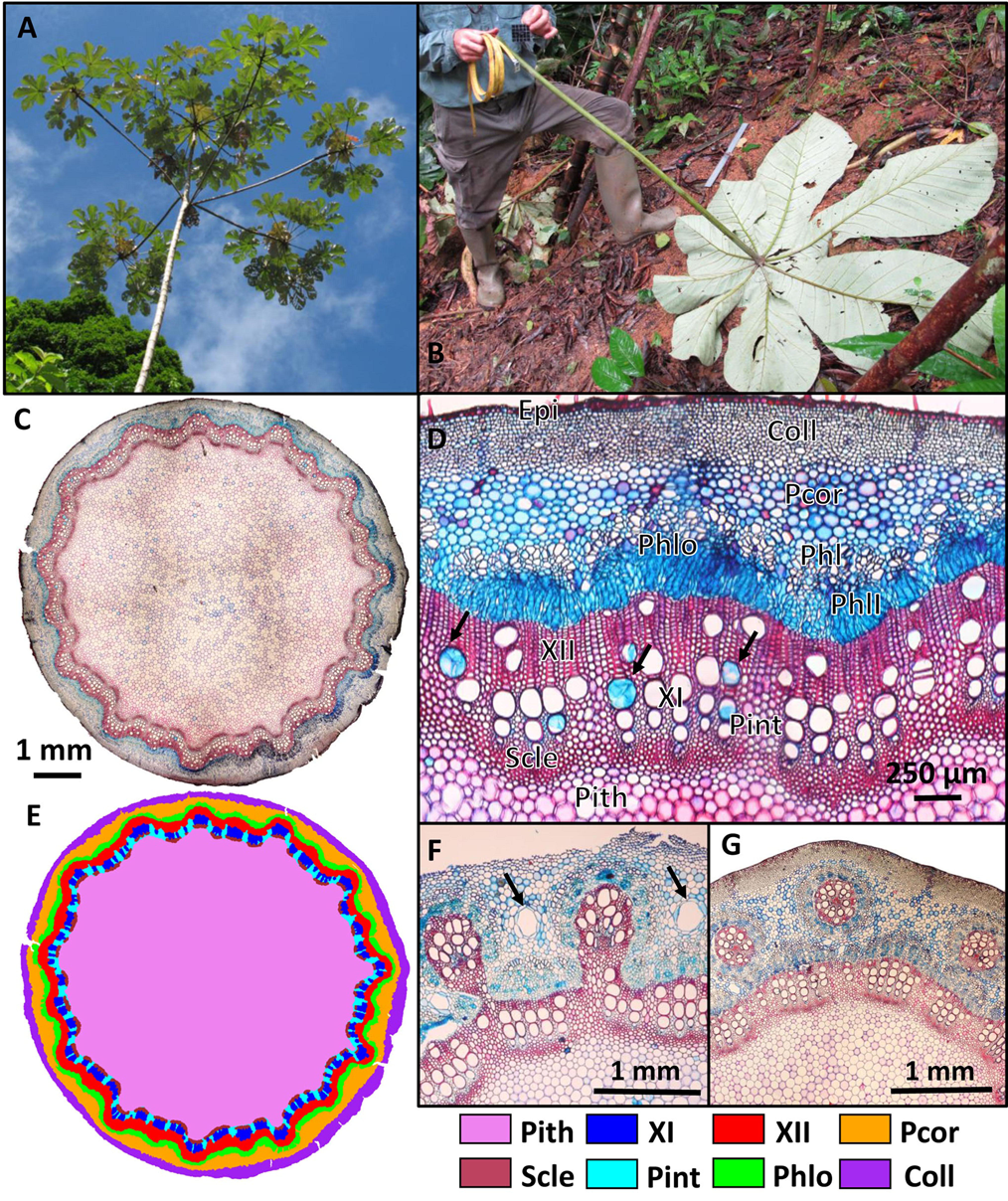
Habits, morphological and petiole-anatomical aspects of C. obtusa (Urticaceae). (A) Habits of C. obtusa. (B) Leaf of C. obtusa. (C) Petiole cross-sectional anatomy of *C. obtusa* in the middle of the petiole. (D) Close-up of petiole constitutive tissues: pith (Pith), sclerenchyma (Scle), interfascicular parenchyma (Pint), primary xylem (XI), secondary xylem (XII), total phloem (Phlo), primary phloem (Phl), secondary phloem (PhII), cortical parenchyma (Pcor), collenchyma (Coll) and epidermis (Epi). Simple arrows indicate tylosed vessels. (E) Tissues and corresponding layer masks studied. (F) “Wavy” cambium with a sub-bicyclic array of vascular bundles. Arrows represent laticiferous canals. (G) Cambial discontinuities, island-like vascular bundles and a bicyclic array of vascular bundles.

Here, we provide a functional assessment of intraspecific LSR in *Cecropia obtusa* by focusing on the lamina-petiole relationship in the megaphyll and its hydraulic implications. First, we report an extensive investigation of the relationships between leaf-level traits to validate the lamina-petiole relationship. We hypothesise that, at this level, correlation properties are comparable to the LSR, thereby enabling further functional assessment. Second, after providing the first description of the anatomy of the petiole, we focus on the lamina-petiole relationship by integrating, according to leaf size (i) the surface areas and proportions of the different component tissues of the petiole, (ii) structural anatomical traits such as the number of xylem vessels, their hydraulic-weighted dimensions and their frequency, and (iii) theoretical hydraulic traits (derived from the Hagen-Poiseuille law) such as hydraulic conductivity and xylem- or leaf-specific hydraulic conductivity.

## MATERIALS AND METHODS

### Study site

This study was conducted in French Guiana along a forestry road in Counami (N5.41430°, W53.17547°, geodesic system WGS84) where the entrance to the road is located 5 km to the east of Iracoubo municipality. The warm wet tropical climate of French Guiana is highly seasonal due to the north-south movement of the Inter-Tropical Convergence Zone. Annual rainfall is 3041 mm year^-1^ and annual mean air temperature is 25.7 °C at Paracou experimental station (Gourlet-Fleury et al. 2004). There is one long dry season lasting from mid-August to mid-November, during which rainfall is < 100 mm month^-1^.

### Plant material, sampling and morphological measurements

In order to explore a wide range of leaf sizes, we studied 25 adult trees (Fig. 1a) and four juvenile trees. Juvenile trees were less than 2.2 m in height and by counting the number of nodes along the trunk (Heuret et al. 2002, Zalamea et al. 2012), their age was estimated at between 8 and 13 months. The dimensions of adult trees ranged from 10.9 to 22.6 m in height, 6.21 to 30.49 cm in diameter at breast height, and the trees were from 7 to 21 years old. Juvenile trees were studied in September 2014. Adult trees were felled regularly from September 2014 to October 2016, between 0800 am and 1030 pm. After felling, all the axes of the crown were tagged according to their position in the tree architecture and all the leaves were tagged according to which axis they belonged, and their rank from the apex. All the leaves in the crown were cut by sectioning the petiole as close as possible to its insertion point on the stem and immediately placed in plastic bags and then in coolers. One to five leaves per tree -hereafter called P-leaves-just fully expanded and positioned on the 3^rd^ or 4^th^ nodes located under any apex, were put aside for the anatomical studies.

To enable a full understanding of the study, here we recall certain plant morphology prerequisites. *Cecropia* leaves are palmatilobate (Fig. 1b and 2). Consequently, there is no single midrib but as many midribs as there are lobes, i.e. a midrib departs from each lobe. We use the term lamina to refer to the flat thin photosynthetic part of the leaf. We use the term petiole to refer to the physical connection between the lamina and the stem. Finally, we use the term leaf to refer to the global structure composed of a petiole and a lamina. In the laboratory, all the leaves were processed at the same time of day to keep them as fresh as possible. The petioles were cut as close to the lamina as possible. The length of the petiole (L_pet_, cm) and two median and orthogonal diameters (mm) of each petiole were measured (Fig. 2a). From these two diameters, a cross-sectional area was derived in the shape of an ellipse (A_pet_, mm^2^; Fig. 2a). For each lamina, the length of the main lobe (i.e. the largest one in the continuation of the petiole; LMlobe, cm) was measured and the number of lobes (Nlobe) counted (Fig. 2b, c). Exactly 2.851-cm2 of lamina in the form of three tablets were sampled with a circular iron punch (diameter = 1.1 cm) taking care to avoid main nerves (Fig. 2d). In the P-leaves, two 1-cm-long petiole segments in the median position were sampled and weighed (Fig. 2e, f). The first segment was used to measure their specific density, expressed as the ratio of dry mass to the fresh volume (PSG: Petiole Specific Gravity; (Williamson and Wiemann 2010); Table 1). The fresh volume of the sample was calculated using an inverse Archimedes principle and a precision balance (CP224S, Sartorius), as the difference between the fresh mass and immersed mass of the sample. Dry mass was derived after drying at 103 °C for three days. The second segment was preserved in 70° alcohol and later used for anatomical measurements (see the section headed ‘Anatomical and hydraulic measurements’).

**Fig 2.**
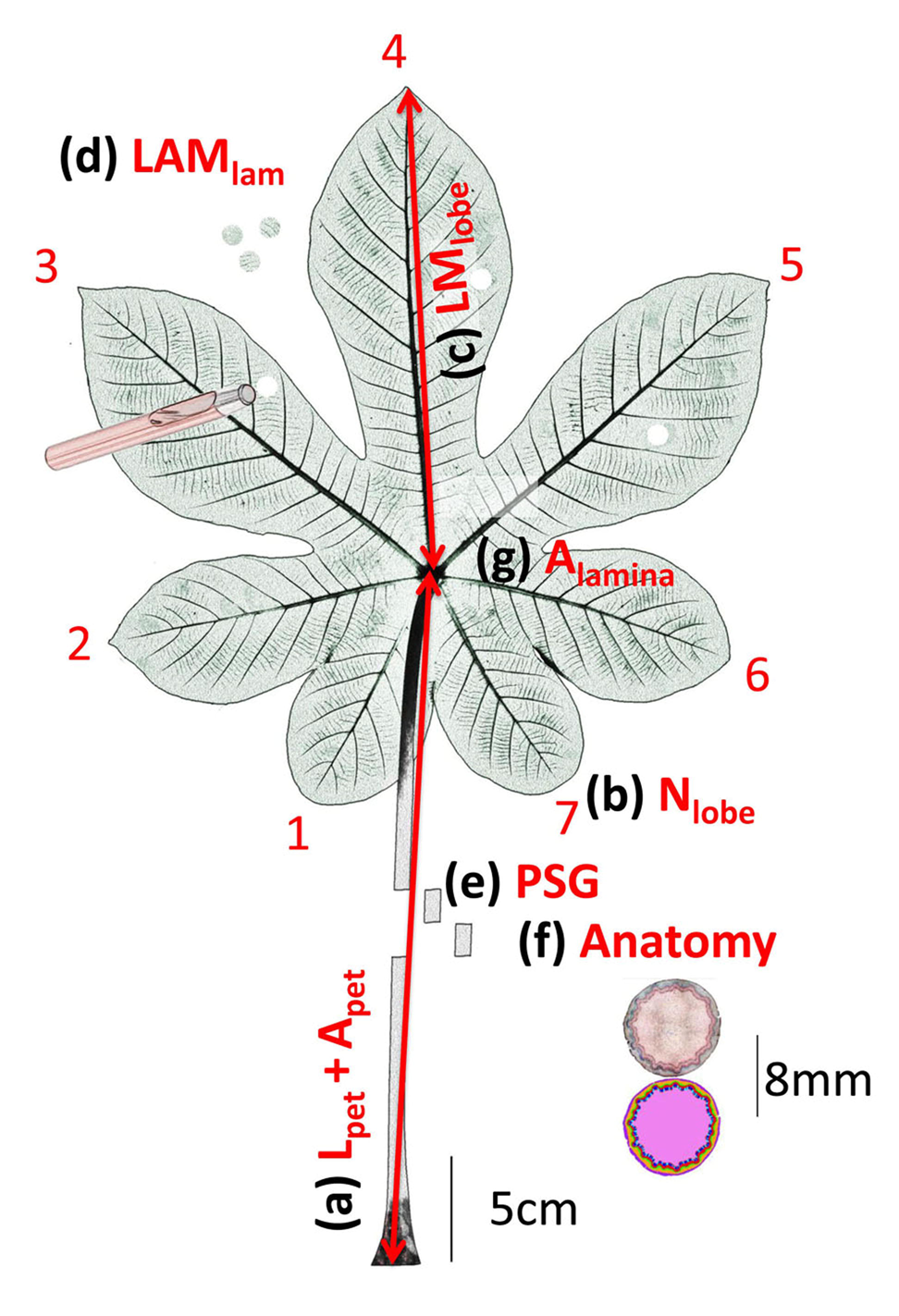
Sampling and measurement design. (a) petiole length (L_pet_); (b) the number of lobes (N_lobe_), here there are 7 lobes; (c) the length of the main lobe (LM_lobe_); (d) The lamina mass area (LMA) is measured based on three tablets if a circular iron punch; (e) in the P-leaves, one segment is used to measurable Se petiole specific gravity (PSG); (f) in the P-leaves, one segment is used for the anatomical study from which is notably derived the petiole cross-sectional area (A_pet_); (g) the lamina surface area (A_lamina_).

**TABLE 1.**
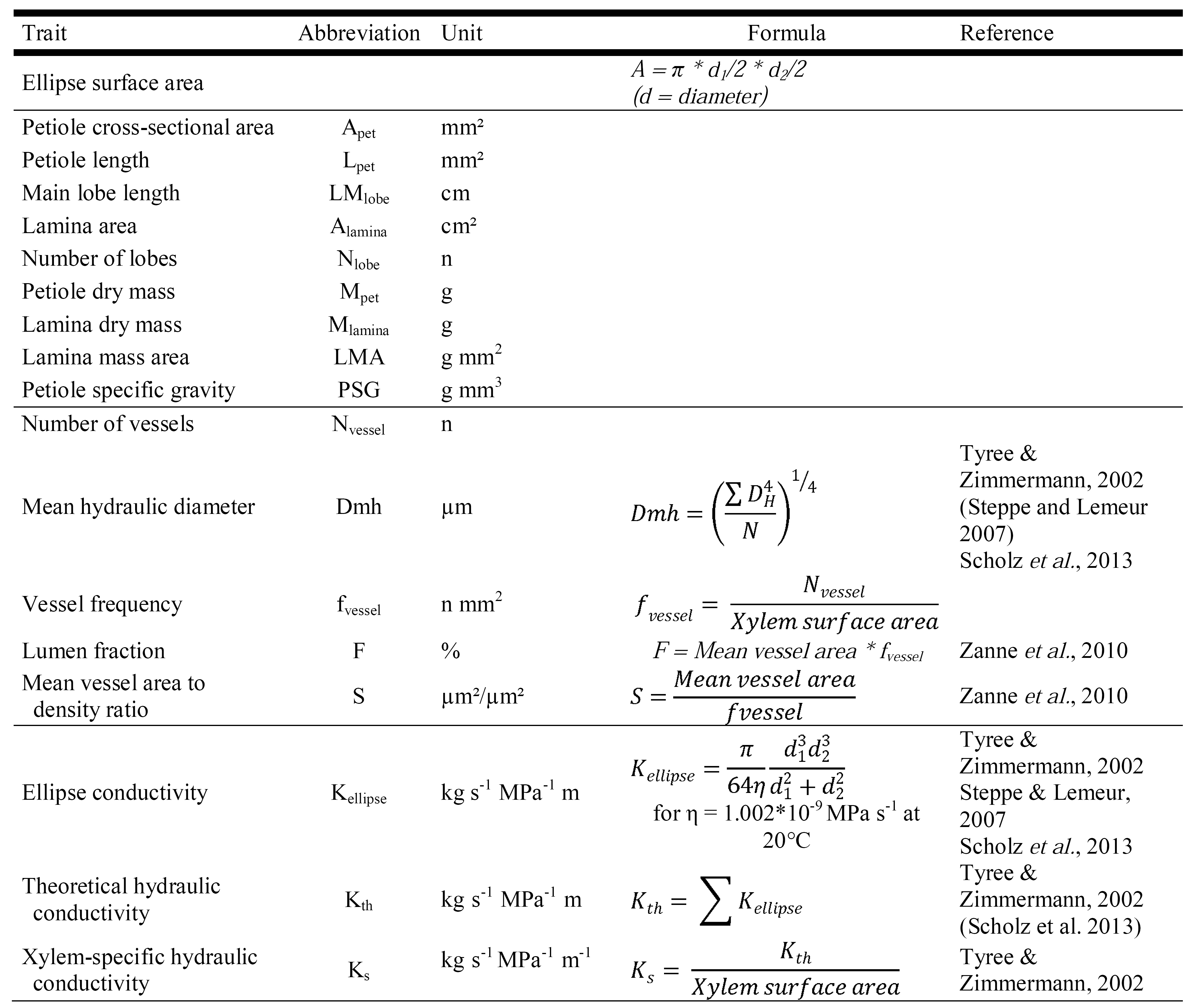
List of the morphological, anatomical and hydraulic traits analysed.

Lamina area (A_lamina_, cm^2^; Fig. 2g) was measured with a planimeter (LiCor 3000A, LiCor Inc., Lincoln, NE, USA). The condition of the lamina (damaged or undamaged) was described. Each leaf (leaf + petiole + 3 tablets) was dried at 70 °C for 72 hours. The dry mass of the petiole (M_pet_, g), the lamina (M_lamina_, g) and the three tablets taken together (g) was measured with a precision scale. The lamina mass area (LMA, g cm^-2^) was derived from the ratio of the dry mass to the known surface area of the three tablets (Fig. 2a).

Data from 2112 mature fully-developed leaves (P and non-P-leaves) were used for morphological validation of the LSR. When we analysed relationships, including lamina area, we only considered leaves with undamaged lamina (*n* = 1072). We also investigated the branching intensity-stem size trade-off (Osada et al. 2015), considering lobes (i.e. main nerves) as homologues of the branching process, and by studying the ratio of the number of lobes to the undamaged lamina area (number of lobes per lamina unit area, we refer to as “lobing” intensity; n mm^-2^) according to the leaf size.

### Anatomical and hydraulic measurements

Because anatomical studies are time consuming, we conducted our study on a subset of 60 P-leaves selected to represent (i) our 29 individual trees to minimise individual effects and (ii) the range including the largest petiole diameters (from 2.59 to 15.41 mm). Anatomical transversal cross-sections, 20 to 50 μm thick, were sampled from the petiole middle segment with a manual microtome (Mikrot L, Schenkung Dapples, Switzerland). All cross-sections were stained in a safranin/astra-blue solution to color unlignified cells blue and lignified cells red.

Measuring anatomy and xylem conductivity in one point is, of course, restrictive to understand the whole vascular architecture and the underlying hydraulic of the entire leaf (Coomes et al. 2008). However, because the aim of this paper is to understand the anatomical patterns underlying the SLR we choose to focus on the median part of the petiole using the petiole-lamina model on megaphylls.

Each cross-section corresponding to a given petiole was digitised with an optical microscope (Olympus BX60; Olympus Corporation; Tokyo; Japan) with x50 magnification and a Canon camera EOS 500D (lens Olympus U-TVI-X; F 0.0; ISO 100; speed 1/25). Three or four close ups were taken of a petiole part at different depths of focus and stacked with the Helicon Focus software (v.6.3.2. Pro, http://heliconsoft.com/). Output pictures were assembled in a panorama using Kolor AutoPanoGiga software (v.3.0.0, http://kolor.com/autopano/) to obtain a complete picture of the cross-section. The digitised cross-sections were processed with CS5 Photoshop software (v.12.0, http://adobe.com/products/photoshop/html). We distinguished eight tissue categories that comprise the petiole anatomy (see anatomical description in the ‘Results’ section). We manually delineated the tissues on the photographs and created layer masks. The masks of these layers were used to calculate the surface area of each tissue and the whole petiole area with the ImageJ software (v.1.43u; http://imagej.nih.gov/ij/). For the xylem, the cropped part of the image in which the vessels were visible was also analysed with the ImageJ software to calculate theoretical xylem hydraulic properties (Abramoff et al. 2004). For each vessel, we calculated its surface area (μm^2^) and its elliptical diameters. These data were used to derive all the anatomical and hydraulic traits listed in Table 1. To study variations in the dimensions of the vessel, we used the mean hydraulic diameter (Dmh, Table 1), i.e. the diameter that all vessels, considered as circles, in a given tissue would have to sustain exactly the same tissue hydraulic conductivity (Tyree and Zimmermann 2002). The number of vessel was counted for primary and secondary xylems and the vessel frequency calculated by dividing this number by the surface area of the related xylem. The lumen fraction (Zanne et al. 2010) refers to the total fraction of empty holes formed by lumens. The ratio of mean vessel area to frequency (Zanne et al. 2010) as a vessel composition index, refers to the relative weight of vessel size versus vessel frequency. Many vessels could be obstructed by tyloses, so we tested for any effect of leaf size on the frequency of obstruction or the effect of obstruction on petiole conductivity. Vessel obstruction was accounted for by removing obstructed vessels from the calculations.

### Data analysis

We used a prediction model to derive estimated lamina area (A_lamina_, cm^2^), since the relationship between the length of the main lobe and the undamaged lamina area is very informative (*R*^2^ = 0.942, *P* < 0.001, Fig. 3). Indeed, laminae used in anatomical studies are sometimes damaged (i.e. biased lamina measurement). We derived the estimated lamina area as:

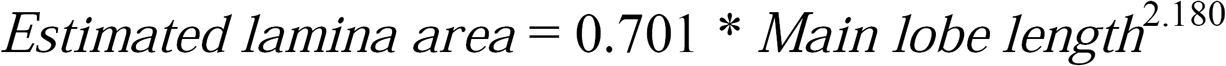

**Fig 3.**
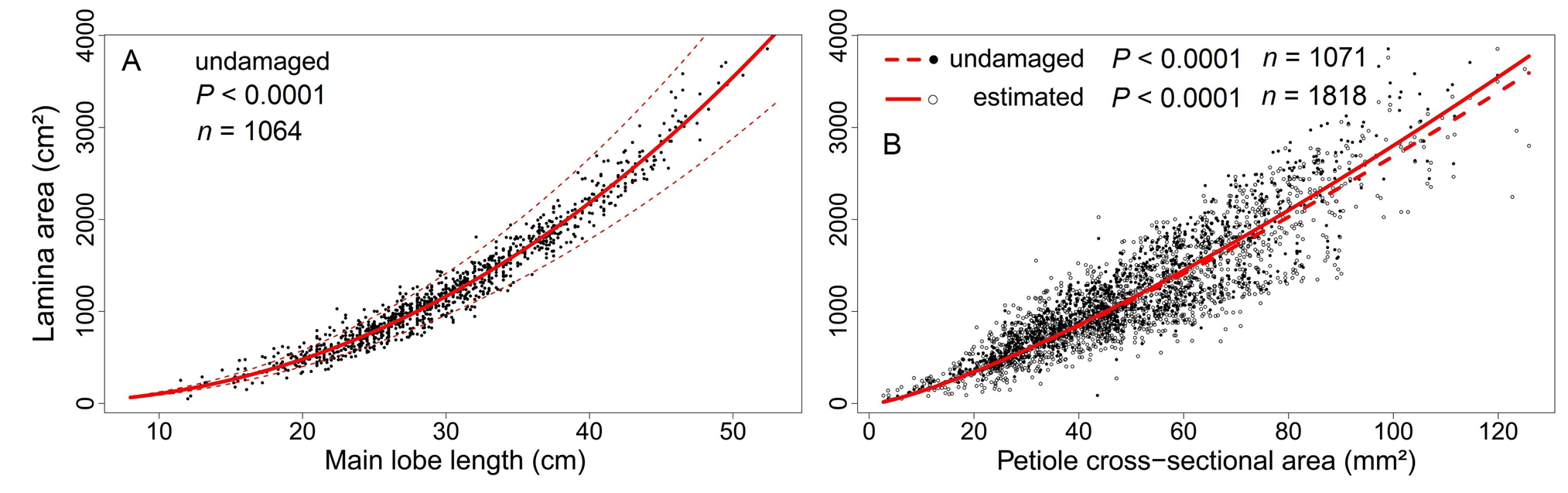
(A) Relationship between the undamaged lamina area (cm^2^) and the length of the main lobe (cm) based on a non-linear least square regression (*P* < 0.001; n = 888). An estimated lamina area can be derived from the length of the main lobe (*y* = 0.90672 * x^2.10950^). Dashed lines represent the confidence interval at 95%. (B) Comparison between undamaged and measured lamina area (filled circles, dashed line; *P* < 0.001; *n* = 892) and estimated lamina area (empty circles, solid line; *P* < 0.001; *n* = 1317) corresponding to the petiole cross-sectional area. Curves are derived from the SMA regression on a log-log scale.

Only this measurement of lamina area was used to compare anatomical and morphological traits.

All statistical analyses were performed with R software (http://CRAN-R-project.org). After log transformation, the relationship between each paired trait was determined with an SMA (Standardized major axis regression; (Warton et al. 2006)), which allows measurement of the error on both the x-axis and y-axis (Harvey and Pagel 1991). These correlation relationships are described as: *y* = b*x*^a^, such as: log(y) = log(b) + a * log(*x*), where *a* is the slope (or allometric exponent) and *b* the intercept (allometric coefficient). A 95% confidence interval was used to decide whether it is significantly correlated or not. A slope test was performed to determine if the slope differed from 1 (H1: a ≠ 1 for an allometric relationship) or not (H0: a = 1 for an isometric relationship). To avoid ambiguity when comparing p-values and decision thresholds, “*P_cor_*” refers to the correlation test, and “*P_slope_*” refers to the slope test (allo- or isometric relationship). SMA were carried out with the (S)MATR package (Falster et al. 2006). The lamina area prediction from the main lobe length was modeled from an NLS (non-linear least squares) with the STATS package.

A paramount *leitmotif* in our study are the scaling relationships between traits, i.e., either iso- or allometric. Indeed, the mathematical relationship has dramatic consequences for data interpretation since it has major functional implications for organism development and phylogenesis (Thompson 1917, Huxley 1932, Niklas 1994). That is why these relationships are briefly examined here. A relationship is expressed as a power law equation, in which the key feature is the scaling exponent characterising the form of the relationship. In biological studies, the relationship generally refers to a size trait on the *x*-axis, whatever the level of organization (cell, tissue, organ, individual, etc.). As the absolute value of the exponent is equal to 1, the relationship is isometric: there is a strictly preserved proportional relationship between the two studied traits independently of their size. In other words, size does not affect the functional nature of the relationship. As the absolute value of the exponent is different to 1, the relationship is allometric: the proportional relationship is not preserved and the y-axis variable fails to keep pace with the x-axis according to the variation in size. In other words, size affects the functional nature of the relationship depending on whether the absolute value of the exponent is greater or less than 1. In the case of a value greater than 1, the y-axis variable increases relatively more rapidly than the *x*-axis variable. In the course of development, the importance of the scaling exponent is such that it can be considered as a trait and a performance condition in its own right since it exhibits its own genetic basis and genetic variations and it is subject to strong heritability (Vasseur et al. 2012).

## RESULTS

### Morphological validation of the lamina-petiole size relationship

Overall, leaf traits were highly and positively correlated (*P* < 0.001; Table 2). There was a significant and positive correlation of allometric form among most traits (A_pet_, L_pet_, A_lamina_, Nlobe, Mpet, Mlamina, LMA; *Pcor* < 0.001; *Pslope* < 0.001; Table 2) except for (i) Alamina which was isometrically correlated with M_lamina_ (*P_slope_* = 0.210) and (ii) LMA which was not correlated with L_pet_ and A_lamin_a (*P_cor_* > 0.05; Table 2). PSG was not correlated with A_pet_, N_lobe_, M_pet_ and M_lamin_a (*P_cor_* > 0.05; Table 2) but was negatively and allometrically correlated with the other measured traits (L_pet_, A_lamina_, N_lobe_; *P_cor_* < 0.05; *P_slope_* < 0.001), except for LMA, which was positively correlated.

The number of lobes per lamina unit area (Nlobe/Alamina) was negatively correlated with Apet (*Pcor* < 0.001; Fig. 4). For a given lamina area, Nlobes increased with a decrease in Apet.

**Fig 4.**
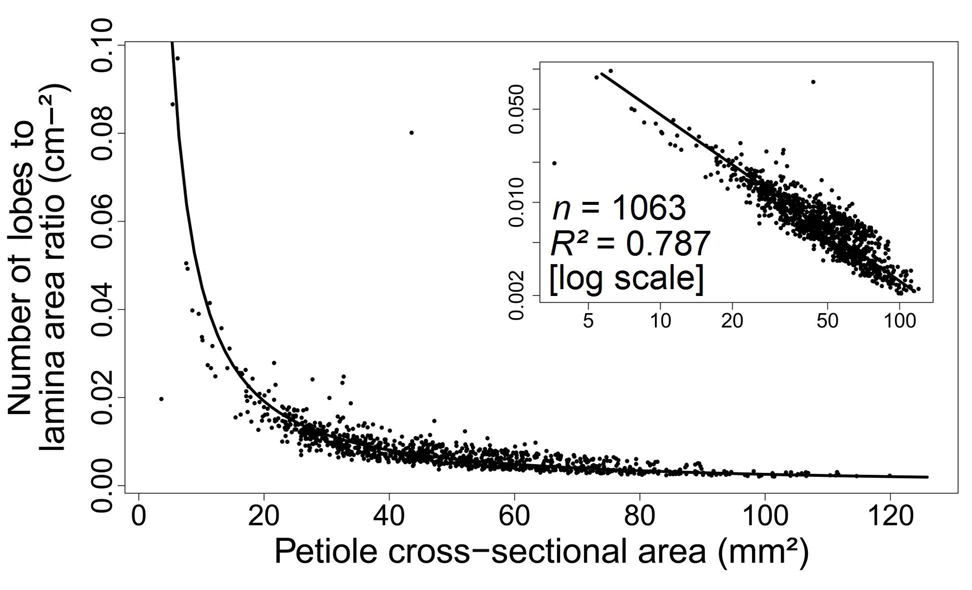
Relationship between the ratio of the number of lobes to undamaged lamina area and the petiole cross-sectional area (*P* < 0.001). The curve is derived from the SMA regression on a log-log scale. Inset shows data on a log-log scale.

### Petiole anatomy

Petioles showed pronounced radial symmetry (Fig. 1c). The central parenchymatous pith formed the main part of the cross-section (Fig. 1c, d, e). The numerous bundles (20 to 80) were radially arranged in the pith periphery, mainly in only one cycle (Fig. 1c), but in a few cases in two cycles (Fig. 1f, g). A cambium gave rise to increments of secondary xylem internally and secondary phloem externally (Fig. 1d). The cambium showed gradual interpetiole diversity with respect to the arrangement of the bundles. From a strictly cyclic structure to a wavy one, and at the extremity, we observed isolated bundles with complete cambium discontinuities in a more cortical position (Fig. 1f, g). Primary and secondary xylem and secondary phloem were easy to identify. Tylosis partially or totally obstructed the xylem vessels (Fig. 1d). A sub-continuous sclerenchymatous shield was present at the interface between the vascular bundles and the pith (Fig. 1d). The vascular bundles were separated by interfascicular parenchyma. Occasional sclerenchyma were present between the secondary phloem and cortical parenchyma. Depending on the extent of secondary growth, the primary phloem was crushed between the secondary phloem and cortical parenchyma. In the most external part, there was a ring of tangential collenchyma under the epidermis, fully contiguous to the cortical parenchyma (Fig. 1d). Laticiferous canals were frequently visible, mainly in the cortex (Fig. 1f) but also in the pith, but were also sometimes completely absent. We distinguished eight component petiole tissues for further anatomical analysis (Fig. 1e): (i) the pith, (ii) the sclerenchymatous shield associated with the vascular bundles, (iii) interfascicular parenchyma, (iv) primary xylem, (v) secondary xylem, (vi) phloem (comprising primary and secondary phloem), (vii) cortical parenchyma and (viii) cortical collenchyma.

### Partitioning of petiole tissue

We now describe the petiole cross-sectional area (A_pet_) both (i) as representative of the variations in several main morphological leaf traits (A_lamina_, M_pet_, M_lamina_, N_lobe_) and (ii) as a proxy of the total leaf size; since there are strong correlations between these traits (Fig. 3; Table 2) and all the anatomical traits are described in relation to the petiole.

**TABLE 2.**
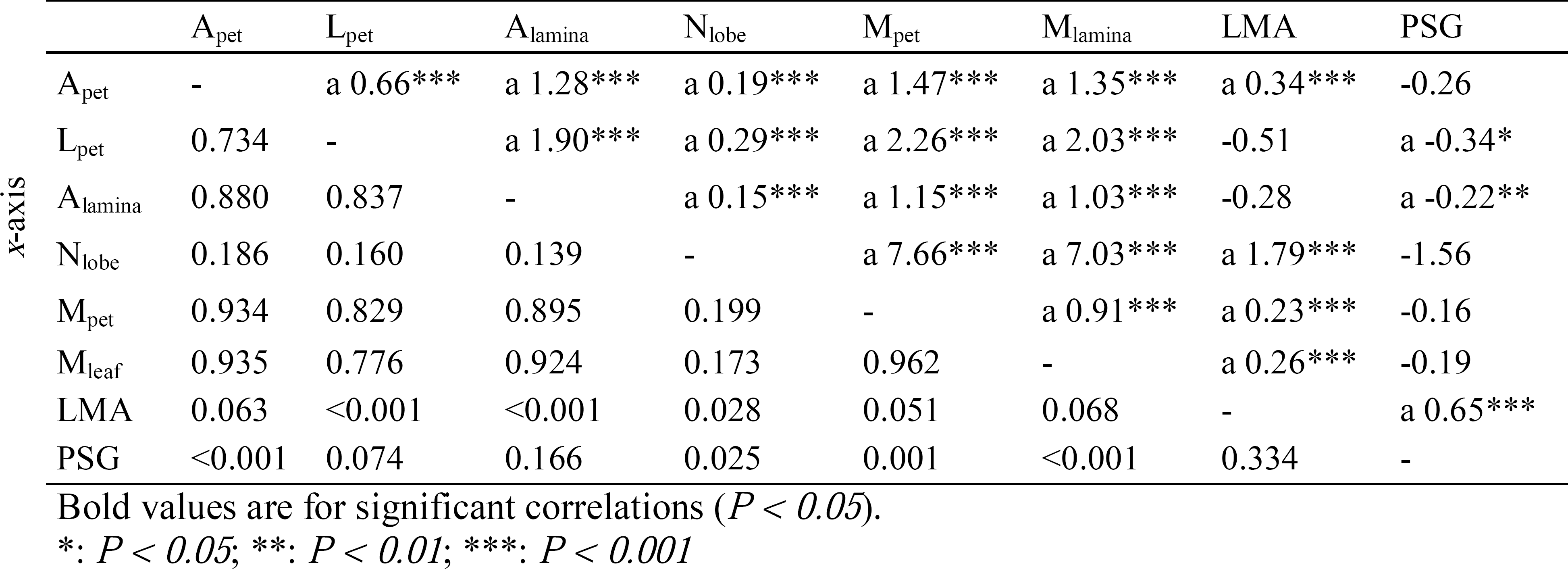
Correlation matrix for leaf-level morphological traits based on SMA regressions on a log-log scale.

All the surface areas of the different tissues observed on the petiole cross-section were significantly and positively correlated with A_pet_ (*P_cor_* < 0.001; Fig. 5a; Supplementary Table 1). Most of the proportions of the different tissues in the petiole cross-section were significantly and positively correlated with Apet (*Pcor* < 0.01; Fig. 5b; Supplementary Table 1), except the primary xylem area, which represented a fixed proportion of Apet (4.89 ± 0.12 %; *Pcor* = 0.337; Fig. 5b,c; Supplementary Table 1). Sclerenchyma, interfascicular parenchyma, primary xylem and secondary xylem represented only small proportions of the petiole cross-section (0.00 to 3.58%; 1.71 to 7.76%; 3.50 to 5.96%; and 2.03 to 9.21%, respectively; Fig. 5b, c; Supplementary Table 1). In contrast, pith, cortical parenchyma, phloem and collenchyma represented significant proportions of the petiole cross-section (31.90 to 59.91%; 8.84 to 29.63%; 2.15 to 17.30%; 8.49 to 23.54%, respectively; Fig. 5b,c; Supplementary Table 1) with high variability depending on the section of the petiole concerned (Fig. 5c; Supplementary Table 1).

**Fig 5.**
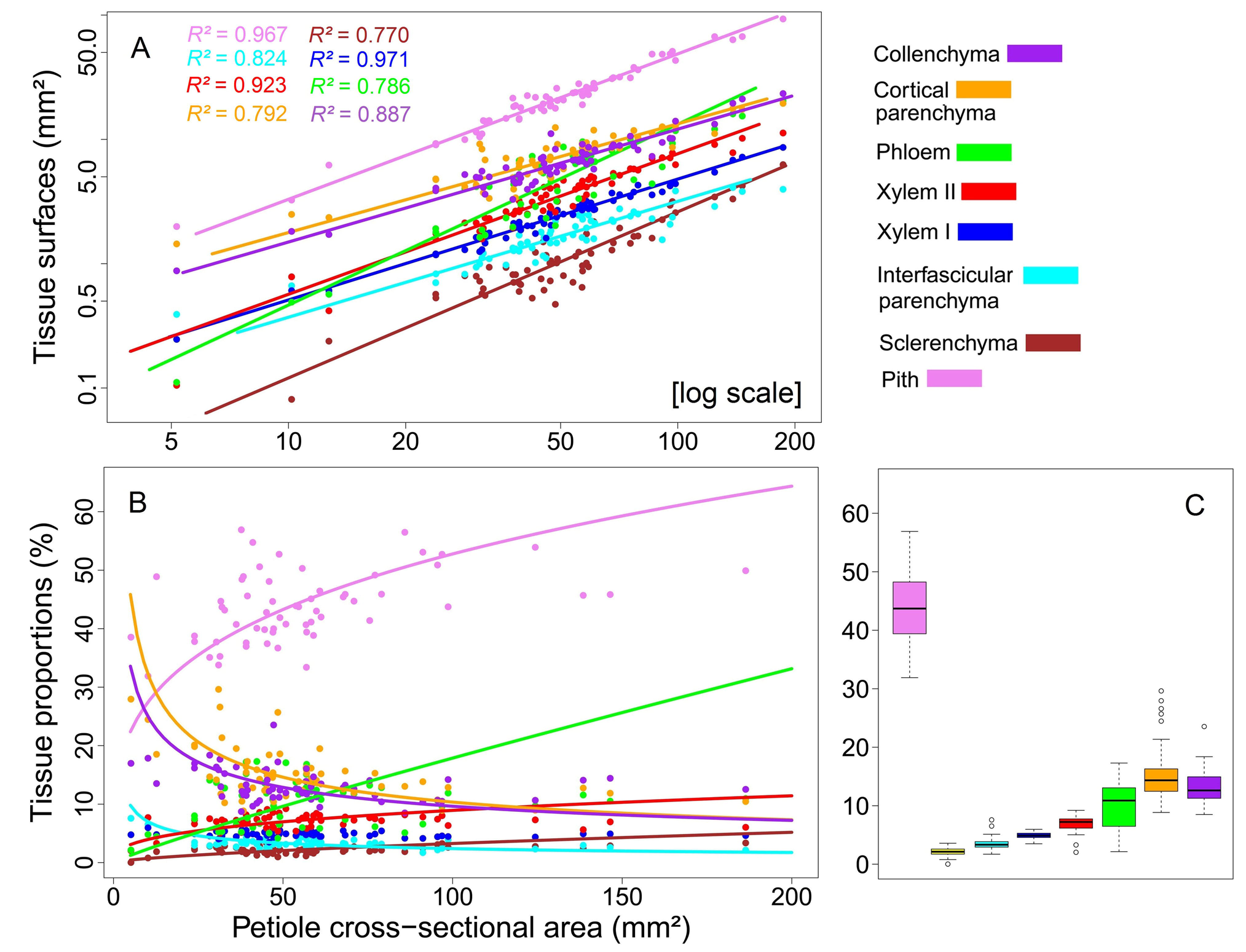
(A) Relationship between the cross-sectional area of the different petiole tissues and the total petiole cross-sectional area on a log-log scale (*P* < 0.001). (B) Relationship between the proportions of the different tissues and the petiole cross-sectional area (*P* < 0.01). Curves are derived from the SMA regression on a log-log scale. (C) Boxplots of the relative proportions of the different tissues according to the petiole cross-sectional area. Pink: pith, brown: sclerenchyma, cyan: interfascicular parenchyma, blue: primary xylem, red: secondary xylem, green: phloem, orange: cortical parenchyma, purple: collenchyma.

A mean of 22.08 ± 1.17% of tissues was calculated for conduction including 4.89 ± 0.12% of the primary xylem, 6.96 ± 0.32% of the secondary xylem and 10.23 ± 0.99% of all the phloem (Fig. 5c; Supplementary Table 1).

### Anatomical and hydraulic considerations of the lamina-petiole size relationship

Hydraulic performances are regulated by three main parameters according to the size of the leaf: surface area of the xylem tissue, the number of vessels (N_vessel_) and the mean hydraulic diameter (Dmh) (Fig. 6). These three traits were closely related (*Pcor* < 0.001), and significantly and positively correlated with Apet (*Pcor* < 0.001; Fig. 6a, b; Supplementary Table 2). However, the frequency of the vessels (fvessel, obtained by dividing the number of vessels by the xylem area; n.mm^-2^) was negatively correlated with A_pet_ (*P_cor_* < 0.05; Fig. 6c; Supplementary Table 2). F was independent of xylem area (*P_cor_* > 0.05; Supplementary Table 2). S was closely correlated with A_pet_ (*P_cor_* < 0.001; Fig. 6d; Supplementary Table 2). Specifically, even for a given fixed vessel frequency, Dmh remained positively correlated with Apet.

**Fig 6.**
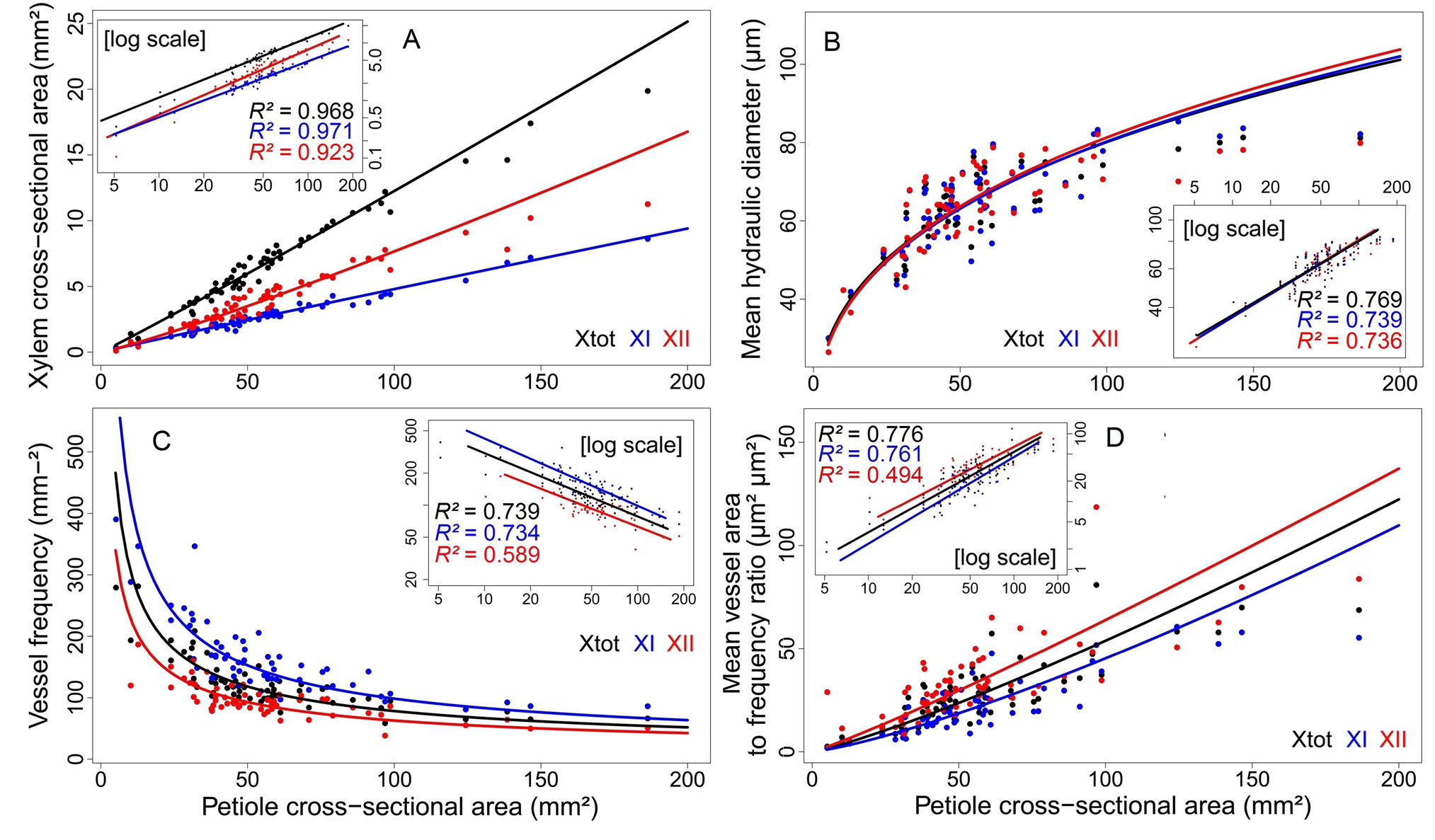
Relationships between the petiole cross-sectional area and (A) the xylem surface area (*P* < 0.001), (B) the mean hydraulic diameter (*P* < 0.001), (C) the vessel frequency (*P* < 0.001), and (D) the ratio of mean vessel area to frequency (*P* < 0.001). Curves are derived from the SMA regression on a log-log scale. Insets show data on a log-log scale. Black: data relative the total xylem. Blue: data relative to the primary xylem. Red: data relative to the secondary xylem.

All these coordinated anatomical adjustments affect theoretical hydraulic properties. There was a positive allometric correlation between the theoretical hydraulic conductivity (K_th_) and A_pet_ with a slope greater than 1 (*P_cor_* < 0.001; *P_slope_* < 0.001; Fig. 7a; Supplementary Table 2). The same was true for A_lamina_ (*P_cor_* < 0.001; Supplementary Table 2). Overall, for each petiole, tyloses did not significantly affect K_th_, whether we took closed vessels into account in our calculation or not, (P = 0.682). There was no correlation between petiole size and the frequency of obstruction (P = 0.636). But there was a significant difference in vessel size between obstructed and unobstructed vessels (P < 0.001) but the opposite pattern was found between primary and secondary xylem. The obstructed vessels were larger in the secondary xylem but smaller in the primary xylem. There was a high positive correlation between Ks and Apet (*Pcor* < 0.001; Fig. 7b; Supplementary Table 2): even with a given fixed xylem area, associated Kth increased with Apet. Kth was also positively correlated with A_lamina_ with a slope greater than 1 (*P_cor_* < 0.001; *P_slope_* < 0.001; Fig 7c; Supplementary Table 2): the larger the leaf, the more supplies go to a given lamina surface area. Moreover, Ks was positively correlated with A_lam_¡_na_ (*P_cor_* < 0.001; Supplementary Table 2).

**Fig 7.**
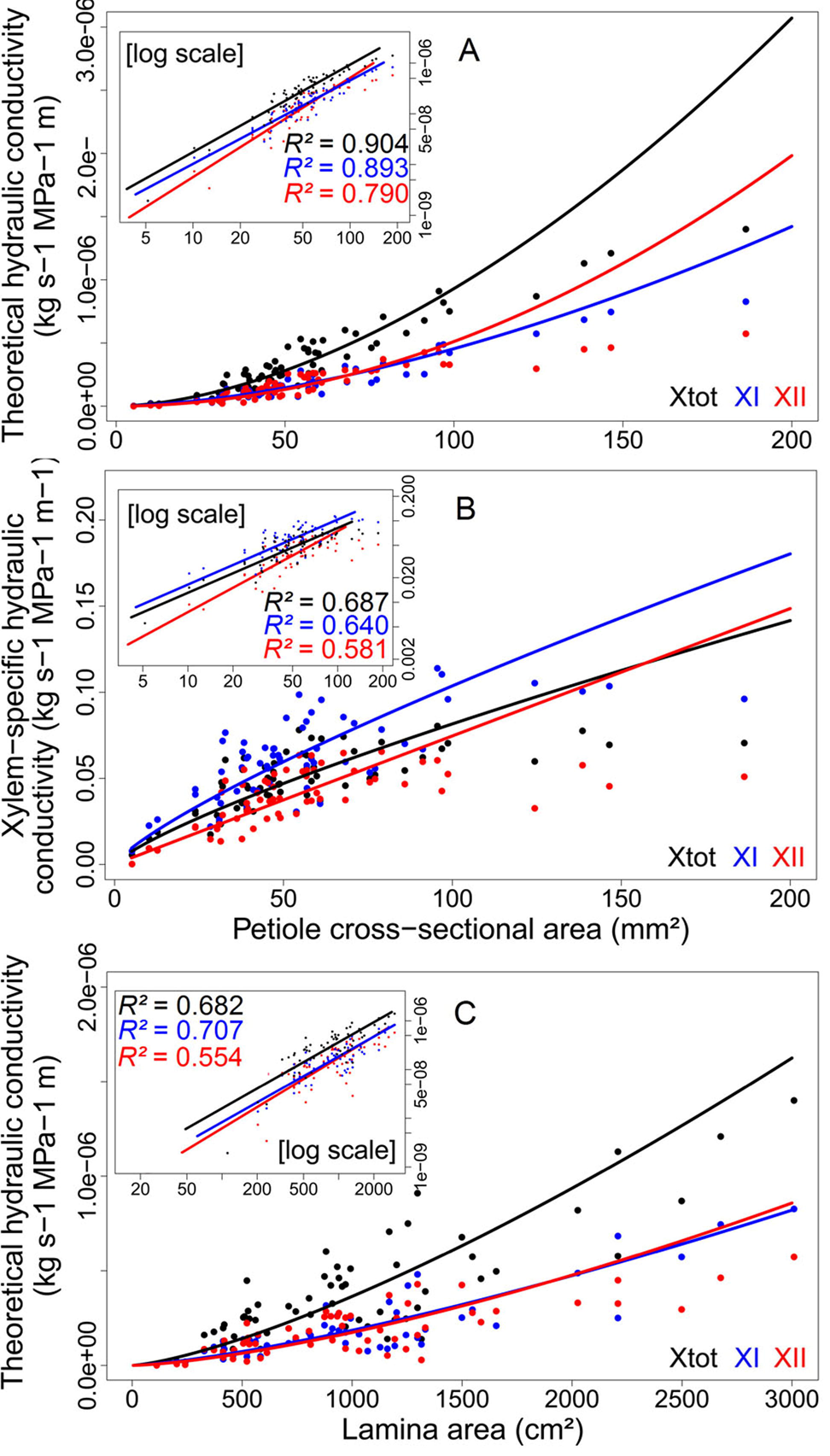
Relationships between the petiole cross-sectional area and (A) theoretical hydraulic conductivity (*P* < 0.001), (B) xylem-specific hydraulic conductivity (*P* < 0.001) and (C) leaf-specific hydraulic conductivity (*P* < 0.001). Curves are derived from the SMA regression on a log-log scale. Insets show data on a log-log scale. Black: data relative to the total xylem. Blue: data relative to the primary xylem. Red: data relative to the secondary xylem.

## DISCUSSION

Our study describes for the first time anatomical structures and mechanisms to better understand the nature of the first premise of Corner’s rule and its underlying hydraulics and mechanical significance. Here, we discuss our three main results: (i) the shift to a leaf-level model for a rigorous functional assessment of the LSR at the intraspecific level, (ii) the pith, not the secondary xylem, as the main driver of variations in petiole size and (iii) and variations in xylem vessel anatomy, not its surface area, as the main determinant of hydraulic requirements.

### Using the leaf model to extend the leaf-stem size relationship

Our results showed strong positive mainly allometric correlations between morphological traits, validating the lamina-petiole relationship as an extensive consideration of the LSR in *C. obtusa.* Most leaf traits were related, not only internode cross-sectional area and lamina area, which are the most frequently studied (White 1983a, 1983b, Brouat et al. 1998, Brouat and McKey 2001, Westoby and Wright 2003, Preston and Ackerly 2003, Sun et al. 2006, Normand et al. 2008, Liu et al. 2010). The allometric form of the relationship between A_lamina_ and A_pet_ should be interpreted with caution, in view of existing results on LSR both at intra- and interspecific levels (White 1983a, 1983b, Brouat et al. 1998, Brouat and McKey 2001, Westoby and Wright 2003, Preston and Ackerly 2003, Sun et al. 2006, Normand et al. 2008, Liu et al. 2010). We argue that confounding intra- and interspecific levels of variation is misleading since the former relates more to developmental processes and mechanisms of adjustment, while the latter relates more to a diversity of strategies, selection pressures and evolutionary pathways. In connection with this distinction, it is interesting to note that although Sinnott’s work (Sinnott 1921) and the first premise of Corner’s rule (Corner 1949) overlap in the global frame of the LSR, Sinnott’s work is more in agreement with an intraspecific and developmental perspective while Corner’s formulations concern an interspecific perspective. The latter assumption is also in agreement with a recent study by (Huang et al. 2016) in which scaling models produced contrasted results between intra- and interspecific data, due to dramatic interspecific variation in stem tissue density and leaf size. Reviewing data produced by White (White 1983a, 1983b), who was the first to test Corner’s rules across species, (Brouat et al. 1998) demonstrated an isometric LSR at the intraspecific level. However, Brouat & McKey (Brouat and McKey 2001) found an allometric form related to the myrmecophytic habit, which is in agreement with our result. Note that even if we focus on the leaf level (lamina-petiole relationship), *C. obtusa* is a hollow-stemmed myrmecophytic tree. As discussed by Brouat & McKey (2001) in connection with the LSR, the allometric form of our lamina-petiole relationship is linked to the pith, which accounts for the largest proportion of the petiole cross-sectional area.

We complete this first point with the branching intensity-stem size trade-off. We found a negative correlation between the number of lobes per lamina unit area and A_pet_. For a given leaf surface, the more numerous the lamina lobes, the thinner the petiole upstream from the lamina. This result is also in agreement with Corner’s rules as interpreted by (Hallé et al. 1978), as “the greater the ramification, the smaller become the branches and their appendages” (Corner 1949). At the branched-complex level, this implies that for both fixed foliar area and photosynthetic rates (Enquist et al. 1999), the degree of branching should be a trade-off with the volume and/or density of the axes. In agreement with this hypothesis, we found a weak but significant trend to decreasing PSG for larger leaves (notably with Alamina, *P* = 0.01; Table 2). This decreasing PSG was largely driven by the proportion of pith tissue (P < 0.001) and underlines the significance of geometry in the support function versus the decrease in PSG in the search for lower volumetric construction costs. In the same line of thought, (Olson et al. 2009) claimed that the negative trade-off between the volume of an axis and its density is a corollary to Corner’s rules and the LSR. This is in agreement with the recent work of (Trueba et al. 2016) at an intraspecific level, since leaf area was shown to be a negative trade-off with both leafing intensity and stem density according to canopy openness in a New-Caledonian woody shrub species. We are first the show that this aspect of Corner’s rules applies to the leaf scale, considering lobes as counterparts of the branching process. Above all, this functional dimension (branching and leafing intensities) is far from being marginally studied (White 1983a, Ackerly and Donoghue 1998, Kleiman and Aarssen 2007, Yang et al. 2008, Tao et al. 2009, Olson et al. 2009, Milla 2009, Whitman and Aarssen 2010, Scott and Aarssen 2012, Dombroskie and Aarssen 2012, Yan et al. 2013, Osada et al. 2015, Huang et al. 2016, Fan et al. 2017), but is far from being integrated in the leaf or global spectra of economy (Westoby et al. 2002, Westoby and Wright 2003, Wright et al. 2004, Díaz et al. 2016), whereas it is directly linked to the optimization strategy of leaf area partitioning (Smith et al. 2017) and hence light-interception performance (Duursma et al. 2012).

Finally, the anatomy of the petiole shows characteristics that are generally associated with stem-level anatomy, like radial symmetry and secondary growth with wood. This is in agreement with the axis-like function of the petiole in the architecture of *Cecropia* (Givnish 1984). In the context of our study, all these features strengthen the homology with a stem-like branched system and the interest of shifting to a leaf model for the rigorous functional and mechanistic assessment of the LSR, i.e. integrating anatomical and hydraulic aspects. However, secondary growth potentially implies an alteration of observed relationships and scaling with time, i.e. with the age of the leaf. This implies that secondary growth continues throughout the lifespan of the leaf, which, however could result in ambiguity with respect to secondary growth, which is generally associated with the renewal of conductive and/or supporting potentials whereas the leaves of *C. obtusa* have a relatively short lifespan (generally < 100 d). So does secondary growth continue throughout the lifespan or only during the phase of primary elongation? This question deserves further investigation in the light of mechanical requirements or for vulnerability to cavitation, for instance.

The three points (i) the positive correlation between A_pet_ and A_lamina_, (ii) the trade-off between leaf size and lobe intensity and (iii) the axis-like anatomy of *Cecropia* petioles, call for a functional analogy between the LSR and the lamina-petiole relationship. This conclusion led us to study the functional significance of anatomical adjustments in the frame of the LSR using a leaf model.

### Pith, not secondary xylem, is the main driver of variations in petiole size

Variations in leaf dimensions could be the result of different processes which take place simultaneously at a macro-anatomical level or at the tissue and cell level. It seems logical that variations in petiole dimensions are primarily driven by variations in tissue surface areas and volumes. In our study, we provided a complete description of petiole anatomy for *C. obtusa* by completing a previous succinct description (Bonsen and Welle 1983). The surface area of all the tissues we studied was found to be positively correlated with A_pet_. However, a surprising result was that the proportions of the petiole tissue were not consistent with the leaf dimensions. Only the proportion of primary xylem to its surface area remained constant. Even if the proportion of variation in the secondary xylem can vary substantially linked to the secondary growth processes, the striking feature is the marked tendency of the variation in the pith to drive variation in the absolute size of the petiole.

Because of a high second moment of areas in flexion of more external supporting tissues (i.e. secondary xylem, sclerenchyma and collenchyma) thanks to the pith, it seems that *Cecropia* petioles have a specific and relatively cheap mechanical strategy for supporting large laminae. The key point is that the proportion of pith is positively correlated with Apet, implying that the second moment of areas of external tissues is positively correlated with petiole size (under the hypothesis of conserved material properties). In other words, changing the proportion of pith with an increase in leaf size allowed for a strong structural effect (from the mechanical point of view) of support tissues (e.g. secondary xylem) as they are pushed centrifugally and since a tube is stiffer than a solid cylinder (for a given materiel and a given amount of this materiel) (Niklas 1992). Here, more research is needed for mechanical quantification. If it holds true, this feature underlines the importance of distinguishing intra- and interspecific levels in scaling models as briefly discussed above. Indeed, here we see how a specific strategy relating to leaf size and form can determine allometric scaling.

### Functional significance of anatomical adjustments by the petiole

The positive correlation we found between the leaf dimensions and K_th_ is in agreement with the results of previous studies showing that larger appendages require larger hydraulic supplies (Preston and Ackerly 2003, Normand et al. 2008, Fan et al. 2017), linked to increasing evaporative demand. However, the allometric form of this relationship suggests that a given lamina surface area for large leaves is supplied with disproportionate amounts of water in comparison with the supply to small leaves. This property could leads to increased evaporation and stomatal conductance but deserves more appropriate ecophysiological measurements. This process is mainly driven by an increase in the vessel diameter (derived from the mean hydraulic diameter, Dmh) rather than an increase in N_vessel_ or in the xylem area. Dmh was positively correlated with Apet whereas fvessel was negatively correlated with Apet. Secondary xylem area and its proportion were positively correlated with Apet, but it is still difficult to disentangle if this process is driven by hydraulic or mechanical needs.

Overall, S was positively correlated with A_pet_, reflecting the greater weight of variations in Dmh compared to f_vessel_ and leading to a larger petiole K_S_ for large leaves. Thus, xylem is hydraulically optimised through variation in Dmh, avoiding a costly and less efficient strategy which would cause more xylem tissues to grow (to support more vessels), and would result in a disproportionately large petiole relative to the lamina. This feature is in agreement with the global interspecific pattern described by Zanne *et al.* (Zanne et al. 2010). However, despite the increasing return on hydraulic performance enabled by the variation in Dmh, there must be a trade-off between Dmh and fvessel. This statement is in agreement with the theoretical model of the packing rule (Sperry et al. 2008, Chen et al. 2012), which predicts that fvessel should isometrically scale with conduit size and thus allows for the area-preserving branching of vessels across different branching levels.

Finally, such variability calls for an investigation of vulnerability to cavitation. Indeed, Scoffoni *et al.* (2016) found that larger vessels in the leaf petiole and major veins are related to vulnerability to cavitation. In our study, we showed that larger leaves have larger vessels suggesting higher vulnerability to cavitation, which would have substantial implications for carbon economy or for dealing with the alternation of dry and wet seasons in our study region or the potential intensification of dry seasons in Amazonia with climate change. Accordingly, we found that tyloses occur preferentially on large vessels in the secondary xylem, while tyloses are usually explained in the light of cavitation mitigation (Micco et al. 2016). Since knowledge is lacking on the intraspecific variability of vulnerability to cavitation and related factors (size, season, ontogeny), further investigations are clearly needed.

### Conclusion

Our study demonstrates that the functional analogy between the LSR and the lamina-petiole relationship holds true based on (i) the positive correlation between Apet and Alamina, (ii) the trade-off between leaf size and lobe intensity and (iii) the described axis-like anatomy of *Cecropia* petioles. From this framework, we point that (i) the pith, not the secondary xylem, is the main driver of variation in petiole size, and (ii) variation in xylem vessel anatomy, not its surface area, is the main determinant of hydraulic requirements by allowing a size-dependent xylem hydraulic efficiency. Our results suggest that hydraulics are a relatively poor explanation as a cause of the Corner’s rule and LSR. Thus, size-related mechanical requirements have to be investigated in the same way, to totally disentangle the frequent but speculative statement that Corner’s rule emerged from hydraulic and mechanical requirements. Such size-dependent xylem efficiency also calls for studying the consequence on stomatal conductance and photosynthesis. Finally, future work will be needed to evaluate if such statement holds true at the interspecific level, by ideally integrating morphological, anatomical and physiological data related to stem, petiole and lamina.

## ACKNOWLEDGEMENTS

The authors thank Julie Bossu, Coffi Belmys Cakpo, Henri Caron, Jocelyn Cazal, Saint-Omer Cazal, Aurélie Cuvelier, Bruno Clair, Aurélie Dourdain, Jean-Yves Goret, Marie Hartwig, Ariane Mirabel, Audin Patient, Pascal Petronelli, Laurent Risser, Dylan Taxile, Valérie Troispoux, Niklas Tysklind and Lore Verryckt for their assistance with field work and measurement of leaf traits. We thank Jacques Beauchêne, Christine Heinz and Nick Rowe for preliminary discussions around the project and first results. We thank Tancrède Alméras for critical and valuable comments on the manuscript.

## FUNDING

S.L. was supported by a doctoral fellowship from CEBA (ref. ANR-10-LABX-0025). This study benefited from an *Investissement d’Avenir grant managed by the Agence Nationale de la Recherche* (CEBA, ref. ANR-10-LABX-0025).

## SUPPLEMENTARY TABLES

SUPPLEMENTARY TABLE 1. Correlation matrix for tissue surface area and proportion traits based on SMA regressions on a log-log scale. Allometric exponents (slopes) are presented.

SUPPLEMENTARY TABLE 2. Correlation matrix between morphological, anatomical and hydraulic traits based on SMA regressions on a log-log scale. Allometric exponent (slopes) are presented.

## Footnotes

1 Manuscript received ———; revision accepted ———

